# A Multiscale Computational Framework Linking Cortical Microtubule Dynamics to Plant Tissue Morphogenesis

**DOI:** 10.64898/2026.06.22.733677

**Authors:** K S Abhijith, Adrija Basak, Orishma Parida, Bandan Chakrabortty

## Abstract

Plant morphogenesis emerges through the coordinated regulation of cell growth and mechanical interactions across multiple spatial scales. A central role in this process is played by cortical microtubule (MT) arrays, which guide cellulose deposition and thereby regulate anisotropic cell expansion. Here, we develop a coupled multiscale computational framework integrating a dynamic vertex model of tissue mechanics with a stochastic model of cortical MT dynamics. Within this framework, MT organization regulates anisotropic cell-wall stiffness, while evolving cell geometry feeds back to influence MT alignment through bidirectional mechanochemical coupling. Simulations show that distinct regimes of MT self-organization generate qualitatively different tissue growth behaviors, ranging from isotropic expansion to strongly anisotropic elongation. Together, our results demonstrate that stochastic a complex coupling of MT self-organization with cell geometry and tissue mechanics is sufficient to generate emergent tissue-scale growth anisotropy, establishing a minimal multiscale framework linking cytoskeletal dynamics to plant tissue morphogenesis.

## Introduction

Unlike animal cells, plant cells are enclosed within rigid cell walls and therefore cannot migrate relative to one another. Consequently, plant morphogenesis emerges primarily through spatially coordinated patterns of cell expansion and division, making tissue growth an inherently mechanical and collective process. A central feature of this process is growth anisotropy, in which cells expand preferentially along specific orientations to generate tissue elongation and organ-level patterning (Baskin, 2005). In plant cells, anisotropic growth is governed largely by the mechanical properties of the cell wall, which are tightly regulated by cortical microtubule (MT) arrays that guide the deposition of cellulose microfibrils (Baskin et al., 1999; Zhao et al., 2020). Because cellulose microfibrils mechanically resist expansion along their orientation, cells preferentially expand perpendicular to the dominant MT alignment (Boudaoud, 2010; Green, 1962). Cortical MT organization therefore acts as a key regulator of directional cell expansion through its control of cell-wall mechanics. Understanding how cortical MT organization regulates growth anisotropy is therefore central to explaining plant morphogenesis.

The primary root of Arabidopsis thaliana provides a well-characterized system for investigating the relationship between cortical MT organization and anisotropic growth (**Figure 1A**). (Hashimoto, 2011). Along the longitudinal root axis, distinct developmental regions exhibit markedly different growth behaviors. Cells within the stem cell niche and meristem undergo limited elongation and display near-isotropic expansion, whereas cells in the elongation zone exhibit rapid and strongly anisotropic growth (Alonso Baez et al., 2026). These contrasting behaviors are closely associated with differences in cortical MT organization. In the meristem, MT arrays are typically weakly ordered and lack a persistent global orientation, while in the elongation zone they become strongly aligned, often oriented transverse to the principal growth axis, thereby promoting directional cell expansion. These spatial transitions suggest that local differences in MT self-organization may play a central role in establishing region-specific growth behaviors during root development. However, despite extensive experimental characterization of these patterns, the mechanistic link between subcellular MT self-organization and emergent tissue-scale growth behavior remains poorly understood. Experimental studies have established strong correlations between MT organization, cell geometry, and anisotropic growth (Bichet et al., 2001; Panteris et al., 2013, 2018). However, disentangling the respective contributions of cytoskeletal organization, wall anisotropy, geometry, and mechanical stress remains challenging because these processes are tightly coupled in living tissues. Moreover, tissue-scale stress distributions and their feedback onto MT alignment are difficult to measure directly in vivo, particularly within developing multicellular structures.

**Figure 1.**
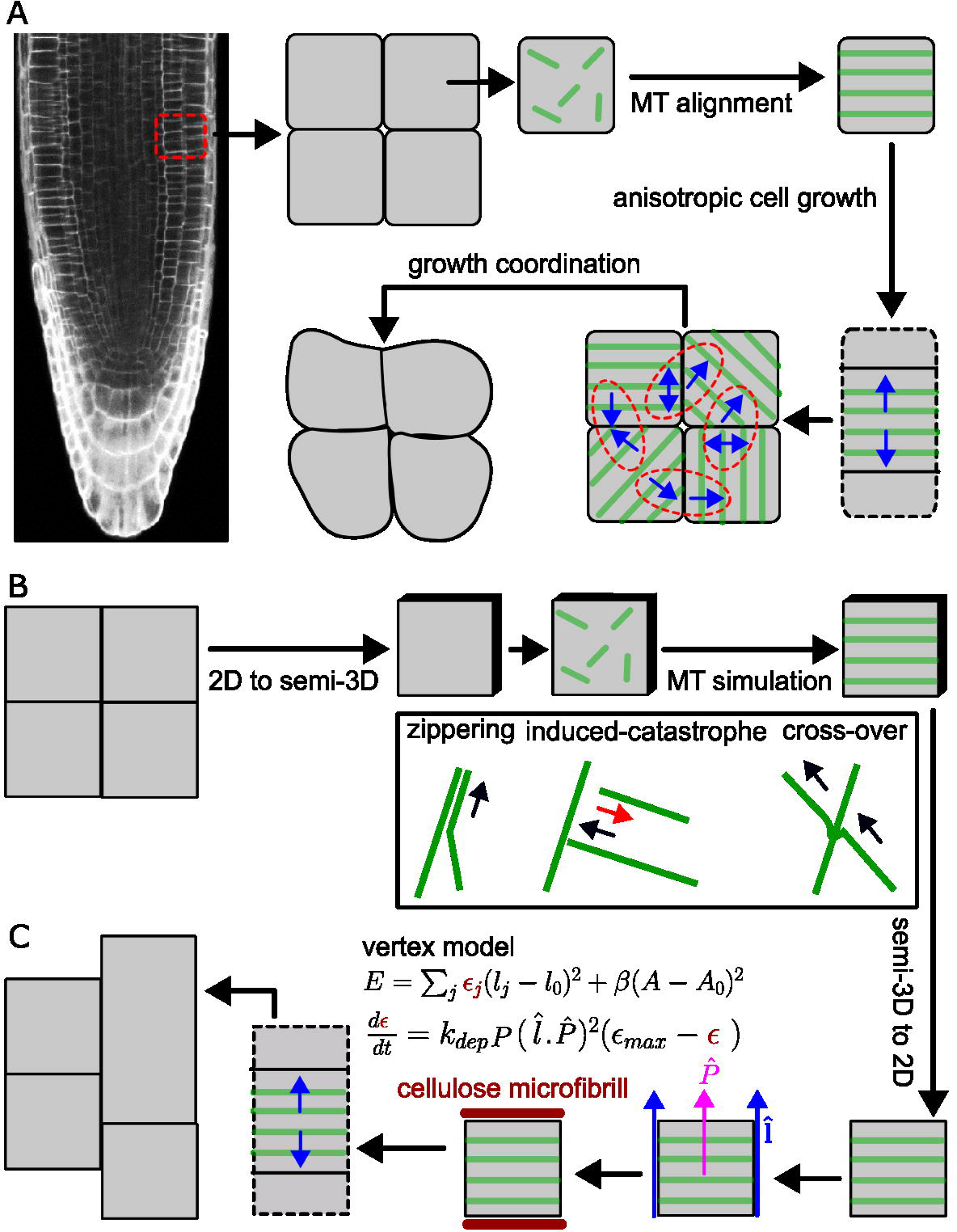
Conceptual and computational framework for microtubule-guided growth coordination in plant tissues. **(A)** Schematic description of MT driven growth coordination in plant tissue: Longitudinal cross-section of an Arabidopsis root tip. A cluster of four cells is highlighted (left, dashed red box). A schematic representation of the cell cluster to explain MT based growth coordination between the cells (right). At the individual-cell level, in the cortex of each cell, nucleation of MTs (green) and their subsequent interactions leads to MT alignment, forming an ordered MT array. Formation of the MT array drives anisotropic cell growth (blue arrow) in the direction perpendicular to the array. At the tissue level, cells can have MT arrays that differ in orientation, resulting in the growth of associated cells in different directions (blue arrow). This, in turn, can cause mechanical conflict (dashed red ellipse) at the interface of each pair of cells. These mechanical conflicts are resolved, i.e., reach mechanical equilibrium, through growth coordination that typically results in cell deformation and, by extension, non-trivial tissue morphology. **(B)** Schematic description of the integrated vertex - MT simulation framework: In our model, tissue consists of cells, which are represented as 2D polygons (quadrilaterals). For simulating MTs using the existing framework that requires 3D cell surfaces (Chakrabortty et al., 2018), for MT simulation on 2D polygonal cell surfaces, we make a copy of the 2D polygonal cell (defined on the x-y plane) and convert it to a semi-3D polyhedral, extruding each cell along the z-direction with a finite thickness (highlighted through a black thick line). On this polyhedral surface, MTs can be simulated using various interaction rules, such as zippering, induced catastrophe, and crossover (Dixit & Cyr, 2004). **(C)** The vector 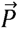 (magenta) is the orientation of the associated MT-array. To simulate cell growth, we implement a dynamic 2D vertex model in which the simulation starts with a tissue configuration of 2D polygonal cells, an existing MT array, and the associated array orientation vector. Cellulose microfibril will be deposited on the cell wall through which MT array is formed. This will make those cell wall stiffer, which is 2D cross-section is equivalent to stiffening (brown) of the complementary (orthogonal) cell walls (edges). This will create anisotropic cell wall stiffness, resulting anisotropic cell growth.

These experimental limitations make computational modeling an essential framework for isolating the individual mechanisms and investigating how local cellular interactions give rise to emergent tissue-scale growth behaviors. Existing computational approaches have provided important insights into either tissue mechanics or MT organization in isolation (Bozorg et al., 2014; Hamant et al., 2008; Huang et al., 2018; Marconi & Wabnik, 2021). Continuum mechanical models have successfully described how tissue geometry and stress distributions influence large-scale organ growth, particularly through finite-element descriptions of tissue mechanics (Hamant et al., 2008). However, because such approaches treat tissues as mechanically continuous media, they do not explicitly resolve individual cell geometries or stochastic MT dynamics. In contrast, vertex-based and other cell-based frameworks represent mechanically interacting cells individually, making them well-suited for investigating how local cellular interactions generate emergent tissue organization (Alt et al., 2017). Separately, stochastic MT models have successfully reproduced the emergence of ordered cortical arrays through dynamic instability, nucleation, and collision-mediated interactions (Chakrabortty et al., 2018; Eren et al., 2015). However, relatively few studies have explicitly integrated stochastic MT self-organization with tissue-scale mechanics within a unified multiscale framework that incorporates feedback between MT alignment and evolving cell geometry (Li et al., 2023). How stochastic MT dynamics at subcellular scales give rise to coordinated growth behaviors across multicellular tissues remains unclear. Likewise, the role of feedback between cell geometry and MT alignment in driving tissue-scale organization is not yet fully understood.

Here, we develop a coupled computational framework that integrates a dynamic vertex model of tissue mechanics with a stochastic model of cortical MT dynamics. Within this framework, MT organization regulates anisotropic cell-wall stiffness, while evolving cell geometry feeds back to influence MT alignment, thereby establishing a bidirectional mechanochemical interaction across scales. To efficiently simulate these multiscale dynamics, we explore the separation of timescales between rapid MT reorganization and slower tissue deformation through a sequential coupling scheme. We then use this framework to investigate how distinct regimes of MT self-organization influence tissue-scale growth behavior. Our simulations show that differences in MT self-organization generate distinct growth regimes ranging from isotropic tissue expansion to strongly anisotropic elongation. More broadly, the results demonstrate that local MT interactions are sufficient to produce emergent tissue-scale growth anisotropy through mechanical feedback, thereby providing a mechanistic link between subcellular MT dynamics and collective tissue morphogenesis. Together, these findings establish a minimal multiscale framework linking stochastic cytoskeletal self-organization to tissue morphogenesis.

## Methods

### Model for simulating tissue growth

To simulate collective cell dynamics within a patch of tissue, we implemented a two-dimensional (2D) dynamic vertex model framework (Alt et al., 2017). This approach enables the coupling of subcellular processes with tissue-scale mechanics while preserving key features of cellular organization. In this framework, cells are represented as polygons defined by vertices, where each vertex may be shared by multiple cells, and neighboring cells share common edges. The mechanical behavior of the tissue is described by an effective energy functional that accounts for edge elasticity, area elasticity, and external confinement. Edge elasticity represents the resistance of shared cell edges to stretching and compression arising from mechanical interactions between neighboring cells. Area elasticity captures the tendency of cells to maintain a preferred size during growth, effectively representing intracellular turgor pressure and resistance to area deformation. Tissue growth and deformation emerge from vertex motion driven by these mechanical interactions, governed by the total energy,

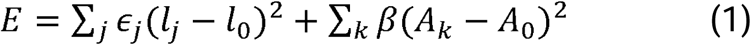

Here, *ε_j_* and *I_j_* denote the stiffness and length of the *j*th edge, *A_k_* is the area of the *k*^th^ cell, and *l*_0_ and *A*_0_ are the preferred edge length and cell area, respectively. The parameter *β* denotes the area elasticity.

For *i^th^* vertex with position coordinate (*x_i_,y_i_*), the corresponding force components (*F^i^_x_*,*F^i^_y_*) are obtained from the energy gradient,

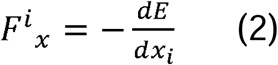

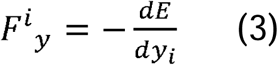

As plant tissue deformation occurs over long timescales and microscopic length scales, tissue dynamics are assumed to be overdamped, with frictional dissipation dominating over inertial effects (Yu et al., 2024). Consequently, vertex acceleration is neglected, and motion is governed by the balance between mechanical forces and viscous drag.

Frictional forces were modeled as linearly proportional to vertex velocities, while hydrodynamic interactions, elastic wave propagation, and thermal fluctuations were ignored. Under these assumptions, vertex motion is governed solely by the balance between mechanical forces derived from the tissue energy functional *E* and dissipative frictional forces characterized by a friction coefficient *Y*, through the following force balance equations,

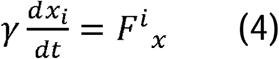

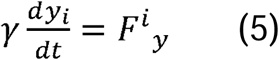

For each time step, the vertex positions are updated (using Eqn. 1 and 2) as follows,

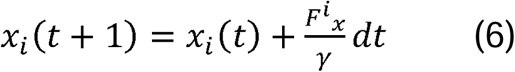

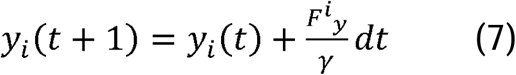

In plant cells, cellulose microfibrils are deposited preferentially along the orientation of cortex-specific microtubules (MTs), commonly referred to as cortical MTs (Paredez et al., 2006). This alignment increases the stiffness of the cell wall along the dominant MT direction (**Figure 1C**). As a result, the anisotropic reinforcement of the wall limits cell expansion parallel to the aligned MT array, leading cells to expand primarily in the direction perpendicular to the dominant MT orientation. These observations highlight the importance of coupling MT dynamics with cell growth to understand the emergence of anisotropic tissue expansion. Furthermore, since cell geometry itself influences MT alignment, this system represents a compelling example of the interplay between cell shape and anisotropic growth.

### Model for simulating MT dynamics

To simulate cortical MTs on the surfaces of 2D cells (polygons) within the vertex model framework (**Figure 1B**), we construct a corresponding thin three-dimensional (3D) representation by extruding each cell along the z-direction with a finite thickness parameter. Because the model focuses on in-plane tissue mechanics, this thickness is chosen to be sufficiently small, thereby preserving the effectively two-dimensional nature of the system while enabling MT simulations on closed cell surfaces. This transformation allows us to use an existing microtubule simulation framework for simulating MT dynamics on the resulting polyhedral geometries (Chakrabortty et al., 2018). In this framework, MT nucleation was assumed to occur isotropically across the cell surface. Individual MTs underwent stochastic dynamic instability, in which growing MTs could transition to a shrinking state through catastrophe events, while shrinking MTs could resume growth through rescue events, governed by prescribed catastrophe and rescue rates. MTs interacted upon collision according to previously established interaction rules, resulting in zippering, induced catastrophe, or crossover events depending on the collision geometry (Dixit & Cyr, 2004). Together, these processes drive the emergence of self-organized cortical MT arrays on the cell surface. Such interaction-mediated ordering represents a hallmark of cortical MT self-organization, whereby local collision rules between individual MTs generate globally aligned arrays without requiring externally imposed directional templates (Lindeboom et al., 2013).

Given the quasi-2D structure of these cells, MTs are expected at steady state to remain largely confined to the x-y plane and exhibit ordered alignment. To quantify this organization, we define an MT-array orientation vector 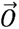, which is perpendicular to the MT-array. Finally, for the original 2D polygonal cells in the vertex model, we define a modified MT-alignment vector 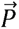 as the projection of 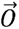 onto the corresponding 2D cell plane,

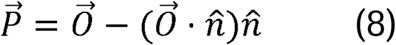

where *n̂* is the unit normal to the associated 2D cell plane. The magnitude 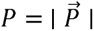 measures the degree of alignment, while the unit vector 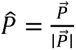 specifies the principal MT-array orientation.

### Integration of the Dynamic Vertex Model and the MT Simulation Model

To implement the bidirectional coupling between MT organization and tissue mechanics, we adopted a sequential update scheme based on the separation of timescales between cortical MT dynamics and plant tissue growth. Since cortical MT reorganization occurs substantially faster than cell growth and tissue deformation, we assumed that MT arrays attain a quasi-steady ordered configuration before appreciable changes in cell geometry occur within the vertex model framework. Accordingly, for each vertex model updated time step (*t*+Δ*t*), MT dynamics were simulated independently on fixed cellular geometries for a duration corresponding to approximately two hour of simulated time, allowing cortical MTs to self-organize and reach steady-state alignment. The resulting MT array orientation vectors 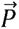 were then used to compute anisotropic cell-wall stiffnesses as follows,

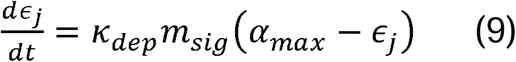

where, *ε_j_* is the stiffness of the *j^th^* cell-edge, *ε*_0_ is the default cell-edge stiffness (i.e., without contribution from MT guided cellulose microfibrils), *ε_max_* is the maximum cell-edge stiffness, *κ_dep_* is the cellulose deposition rate, and contribution from MT guided cell-edge stiffness to the cellulose deposition is incorporated via,

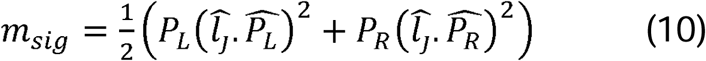

where, 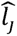 denotes a unit vector along the *j^th^* cell-edge, *P_L_* and 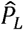 represent the magnitude and direction of the MT alignment vector associated with the cell on the left side of the edge, and *P_R_* and 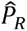 represent the magnitude and direction of the MT alignment vector associated with the cell on the right side of the edge.

The updated edge stiffnesses were subsequently incorporated into the tissue energy functional to compute vertex forces, and accordingly update vertex positions (see Eqn. 6 and 7) within the vertex model framework. The resulting deformed cellular geometries were then used as inputs for the next MT simulation cycle, thereby establishing a feedback loop between MT organization and tissue-scale mechanical deformation. Through this iterative scheme, local MT alignment influences tissue deformation via anisotropic wall mechanics, while mechanically induced changes in cell geometry subsequently alter MT organization, thereby generating a multiscale mechanochemical feedback system. This bidirectional coupling enables the emergence of anisotropic growth through the interplay between MT dynamics and tissue mechanics.

### Simulation parameterization

For vertex model simulations, to make it nondimensionalized, the reference cell-wall stiffness *ε*_0_ and reference cell area *A*_0_ are set to unity, i.e., *ε*_0_ = 1 and *A*_0_ = 1. In the initial configuration (*t* = 0), cells are assumed to be square-shaped with an area *A_ini_* equal to half the reference area, i.e., *A_ini_* = 1/2. Consistent with this, the rest length of the cell edges is set to 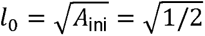. Due to the elastic nature of the forces, cells grow naturally until their area approaches the reference value, *A*_final_(*t* = *t*_max_) ≈ *A*_0_, where *t*_max_ is the maximum simulation time. Cells are assumed to be incompressible, corresponding to the limit β → ∞ (implemented as a sufficiently large value in simulations). The maximum cell-edge stiffness *α_max_* is set to a very large vale to account for strong anisotropic reinforcement, persistent mechanical memory, more resistant cell walls. The friction coefficient *γ* primarily controls the timescale of vertex relaxation rather than the qualitative mechanical behavior of the tissue. Accordingly, we set *γ* to a value that ensures numerically stable dynamics while maintaining the overdamped regime. We verified that, for sufficiently large values of *γ*, the qualitative behavior remains unchanged, confirming that this parameter primarily rescales the relaxation timescale. The remaining parameter, the cellulose deposition rate *κ_dep_* is treated as a free parameter that controls the rate of cell-wall reinforcement in response to microtubule-guided cellulose deposition. For the MT simulations, realistic plant cells typically possess surface areas on the order of ∼ 1000 *µm*^2^. However, in our the nondimensionalized vertex model framework maximum area of a cell is set to *A*_0_ = 1. Since the MT simulation framework primarily depends on the relative density of MT nucleation events rather than the absolute physical cell size, accordingly we rescaled MT dynamic parameters involving spatial or geometric scales (**Table 1**). These include the MT plus-end growth velocity (*ν*_+_), minus-end shrinkage velocity (*ν*_-_), treadmilling velocity (*ν^tm^*), and the MT nucleation rate.

**Table 1:** MT Parameters:

| Parameter | Dixit & Cyr, 2004 | Our Model |
| --- | --- | --- |
| $v_+$ | $0.08 \mu\text{M sec}^{-1}$ | $0.004 \mu\text{M sec}^{-1}$ |
| $v_-$ | $-0.16 \mu\text{M sec}^{-1}$ | $-0.007 \mu\text{M sec}^{-1}$ |
| $v^{tm}$ | $0.01 \mu\text{M sec}^{-1}$ | $0.0004 \mu\text{M sec}^{-1}$ |
| $kNuc$ | $0.01 \mu\text{M}^{-2} \text{sec}^{-1}$ | $5.0 \mu\text{M}^{-2} \text{sec}^{-1}$ |

**VM parameters:**
| Parameter | Values |
| --- | --- |
| $dt$ | 0.00001 |
| $A_0$ | 1.0 |
| $\gamma$ | 1.0 |
| $\beta$ | 1000.0 |
| $\epsilon_0$ | 1.0 |
| $\epsilon_{\max}$ | 100000 |
| $kDep$ | 10.0 |

## Results

### Simulation Validation

To validate the implementation of the vertex model framework, we first examined the relaxation dynamics of a mechanically compressed cell initialized with an area smaller than its preferred area. Starting from (*A_ini_* = 1/2), the cell progressively expanded toward its preferred equilibrium area, (*A*_0_ = 1). Concurrently, the total mechanical energy decreased monotonically and eventually reached a steady-state plateau, indicating relaxation toward mechanical equilibrium (**Figure 2A**). Next, we investigated whether the spatially rescaled microtubule (MT) simulation framework preserved the characteristic self-organization dynamics of cortical MT arrays. To this end, simulations were performed using both a realistic cell geometry and the reduced cell geometry employed in our framework. In both cases, the degree of MT alignment increased progressively with time (**Figure 2B**) before reaching a steady-state plateau corresponding to the formation of stable ordered MT arrays. Furthermore, both geometries exhibited comparable transitions from initially disordered MT configurations to globally aligned cortical MT arrays, demonstrating that the reduced geometry preserves the essential MT self-organization dynamics. To verify the implementation of MT-guided cell-edge stiffening, we simulated MT dynamics in a single cell using the default MT-MT interaction rules and dynamic instability parameters. Under these conditions, the MT array remained unbiased, resulting in a spatially uniform increase in cell-edge stiffness over time (**Figure 2C**). This behavior is expected because, in the absence of directional bias arising either from MT dynamics, MT interactions, or cell geometry (square cell), cellulose deposition occurs uniformly along all cell edges. To determine whether biased MT dynamics could generate anisotropic cell-edge reinforcement, we introduced elevated catastrophe rates at selected cell edges (Ambrose et al., 2011). This perturbation promoted the preferential accumulation and alignment of MTs along the remaining cell edges.

**Figure 2.**
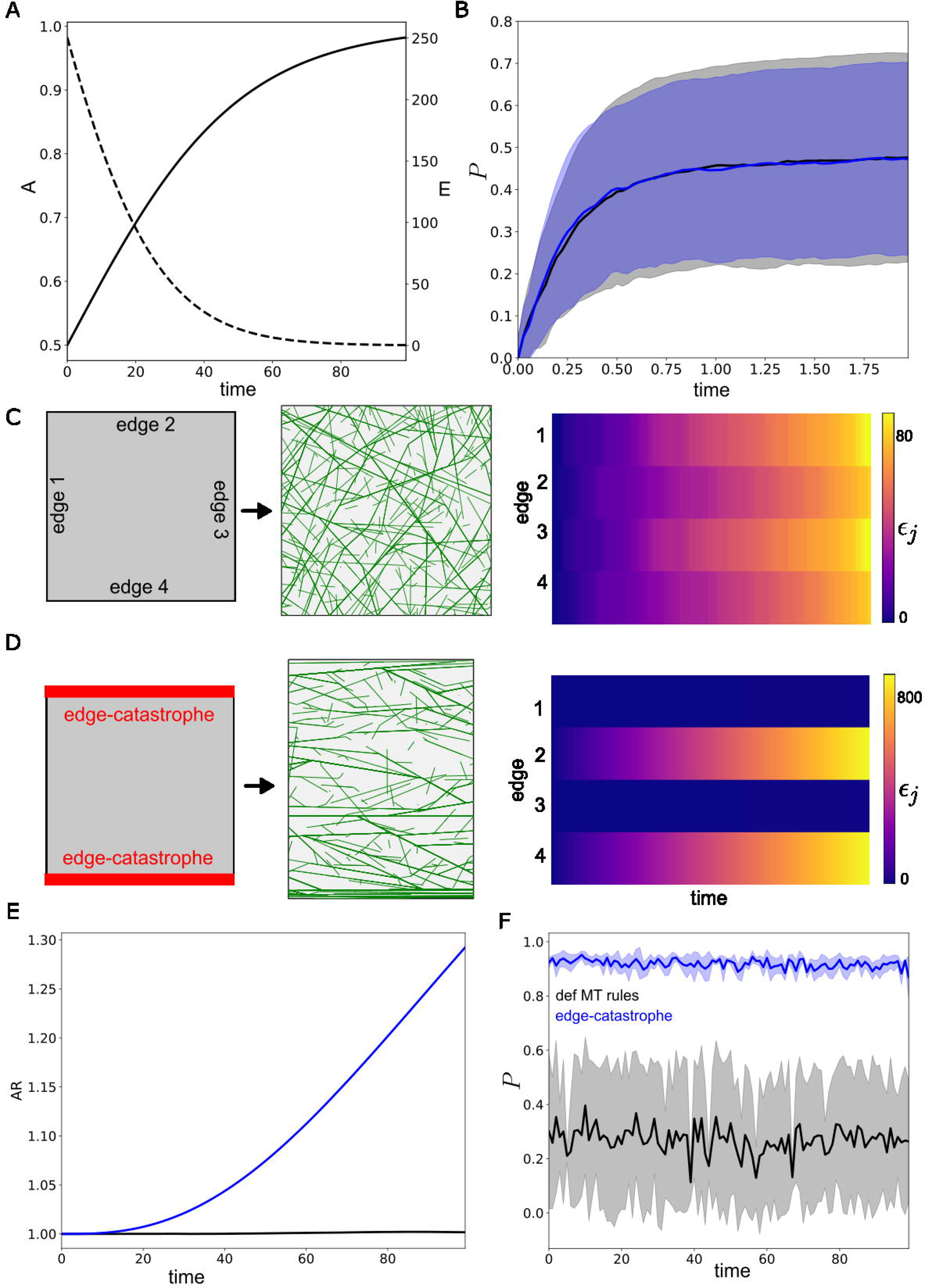
Validation of the integrated vertex model–microtubule simulation framework. **(A)** Validation of the vertex model: Growth simulation of an initially rectangular cell of area *A_ini_* = 0.5 without MT simulation. Plot of the time evolution of the cell is A (black solid line) and cell energy *E* (black dashed line). **(B)** Validation of the MT simulation for the cell size rescaled to the maximum vertex model cell area surface is *A*_0_ = 1.0, compared to the actual plant root cell, having a typical surface area ∼ 1000 µm^2^. Comparative plot of the time evolution of the steady state value of the MT order parameter (P), when simulated using a cell of surface area *A*_0_ = 1.0 (black line) and a cell of surface area *A*_0_ = 1000 µm^2^ (blue line). In this plot, time is in unit of hour. **(C)** Validation of the integrated vertex model and MT simulation framework: Simulated configuration of MTs on a rectangular cell surface, where MTs are simulated using the default interaction rules (as described in Figure 1B) **(D)** Simulated configuration of MTs on a rectangular cell surface, where MTs are simulated using the default interaction rules along with additional catastrophe at selected cell edges (red). The magenta line shows the orientation of the MT array (middle). Heat map, showing the time evolution of the wall stiffness at the cell edges (right). **(E)** Comparative plot of the time evolution of the aspect ratio of the cell, when simulated using only default MT-MT interaction rules (black), and MT-MT interaction rules along with edge-catastrophe (blue). **(F)** Comparative plot of the time evolution of the steady state value of the MT order parameter *P*, when simulated using only default MT-MT interaction rules (black line), and with MT-MT interaction rules along with edge-catastrophe (blue line). For vertex model, time unit is nondimensional with step, dt = 0.00001. For details on parameters please refer to **Table 1**.

Consequently, cell edges parallel to the dominant MT array exhibited higher stiffness values that increased progressively over time (**Figure 2D**), leading to anisotropic mechanical reinforcement. Finally, we quantified the morphological consequences of anisotropic cell-edge stiffening by fitting an ellipse to the final cell shape and calculating its aspect ratio (AR). Cells exhibiting anisotropic edge stiffness developed more elongated morphologies and displayed significantly higher AR values than cells with isotropic stiffness (**Figure 2E**), confirming that MT-guided mechanical reinforcement can drive anisotropic cell shape formation. We also examined the degree of MT alignment in each case, which showed edge catastrophe-based simulation had higher degree of MT alignment, consistent with the absence of coherent alignment in default MT-rule based simulations (**Figure 2F**).

### MT self-organization drives anisotropic cell growth

Having validated the model, we next investigated how MT orientation and the resulting memory of cellulose microfibril deposition influence cell morphology. Previous studies have shown that following cell division, a fraction of MTs re-nucleate at angles closely aligned with the orientation of the pre-division MT array, suggesting the existence of a structural memory mechanism (Lindeboom et al., 2013)Motivated by this observation, we simulated the growth of a single cell with initially isotropic MT orientations (**Figure 3A**). Under these conditions, MT organization remained unbiased, resulting in isotropic cell growth and a low, spatially uniform pattern of cell-wall stiffness. We then incorporated MT memory into the simulation to assess its effect on cell shape anisotropy. Specifically, approximately 30% of newly nucleated MTs were biased toward the average resultant MT array orientation vector (P) of the preceding time frame. Across multiple simulation runs, the inclusion of MT memory consistently generated anisotropic growth, oriented either longitudinally or transversely relative to the dominant MT alignment (**Figure 3B**). These results indicate that the final orientation of the MT array, and consequently the direction of anisotropic growth, is strongly influenced by the initial MT nucleation orientation and its subsequent propagation through memory. Because plant cell behavior is regulated not only by intrinsic factors but also by external mechanical cues (Hamant & Haswell, 2017), we next examined whether MT memory could be guided by an external directional signal, such as mechanical stress. To this end, we introduced a bias that promoted MT nucleation along a prescribed external cue direction. Under these conditions, MT arrays progressively aligned with the imposed cue, leading to anisotropic cell growth in the direction perpendicular to MT orientation (**Figure 3C)**. Thus, external cues can effectively steer MT organization through memory-driven reinforcement, thereby controlling the direction of cell growth. To quantify the influence of MT memory on cell morphology, we compared the aspect ratio (AR) of cells across the different simulation conditions. Cells lacking MT memory exhibited only minor changes in AR and remained nearly isotropic throughout growth. In contrast, incorporating MT memory resulted in a substantial increase in cell shape anisotropy, with the final AR increasing by approximately 25% relative to the no-memory condition (**Figure 3D**). These findings demonstrate that MT memory is sufficient to generate and maintain anisotropic cell growth, providing a mechanism by which transient MT organization can be translated into persistent morphological changes.

**Figure 3.**
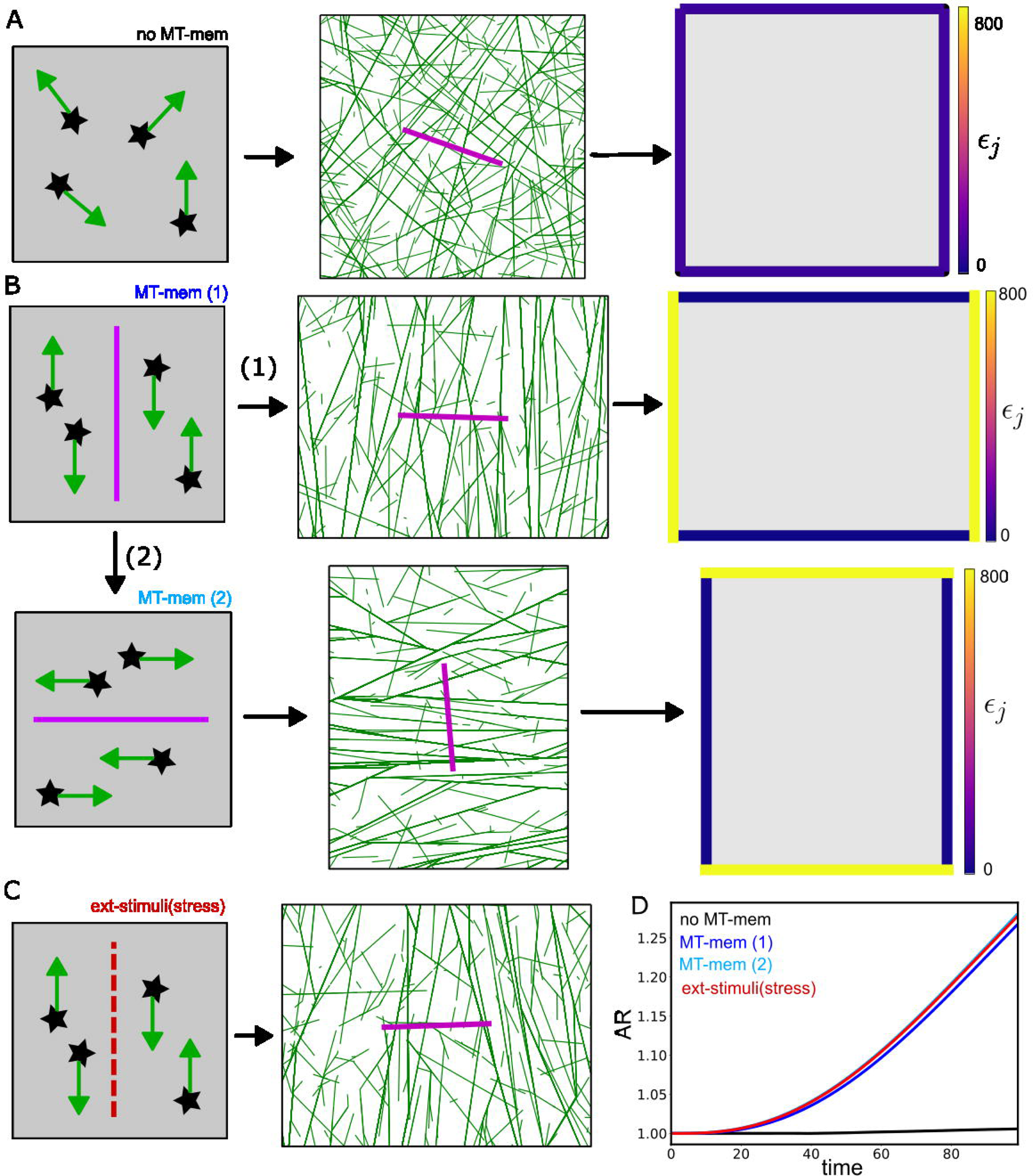
MT memory and external stimulus-based nucleation shape microtubule organization and cell wall mechanics. **(A)** Simulation using MT memory-based nucleation, where MTs are simulated using the default interaction rules (as described in Figure 1B), Heat map, showing wall stiffness at the cell edges (right). **(B)** Schematic description of the implementation of pre-seeded MT nucleation that depends on the existing MT array orientation. The MT array orientation at time t is computed, which is used as a reference for a fraction (f) of nucleation to happen along this direction, at time t+1 (MT memory-based nucleation). The remaining fraction (1-f) happens through isotropic nucleation. Simulation using MT memory-based nucleation, where MTs are simulated using the default interaction rules (as described in Figure 1B), along with the MT memory-based nucleation, which gave two possible configurations: (1) longitudinal cell expansion, and (2) transverse cell expansion. The magenta lines show the orientation of the MT array (middle). Heat map, showing wall stiffness at the cell edges (right). **(C)** Schematic description of the implementation of pre-seeded MT nucleation that depends on the external stimuli, e.g., anisotropic mechanical stress (left). At any time, the direction of the stress is used as a reference for a fraction (f) of nucleation to happen along its direction (external-stimuli-based nucleation). The remaining fraction (1-f) happens through isotropic nucleation. Simulation using external-stimuli-based nucleation: Simulated configuration of MTs on the individual cell surfaces (middle), where MTs are simulated using the default interaction rules (as described in Figure 1B) along with the external stimulus-based nucleation. The magenta lines show the orientation of the MT array. **(D)** Comparative plot of the aspect ratio of the cell, when simulated using only default MT-MT interaction rules (black, refer to Figure 1B), MT-MT interaction rules along with MT memory-based nucleation (blue), and MT-MT interaction rules along with external-stimuli-based nucleation (green). For vertex model, time unit is nondimensional with step, dt = 0.00001. For details on parameters please refer to **Table 1**.

### MT self-organization drives anisotropic tissue growth

We next sought to determine how the mechanical behavior observed at the single-cell level translates to the tissue scale. Unlike isolated cells, cells within a tissue must remain mechanically coupled to their neighbors while growing, raising the question of how tissue-scale morphology emerges from local MT-guided reinforcement under these constraints. To address this, we constructed a tissue composed of a square grid of cells and simulated three conditions: (i) no MT memory, (ii) MT memory derived from pre-existing MT/cellulose patterns, and (iii) MT memory guided by an external directional cue. Throughout the simulations, we monitored tissue morphology and the evolution of cell-wall stiffness along cell edges. As expected, tissues lacking MT memory showed no significant changes in cell-wall stiffness, and tissue growth remained isotropic throughout the simulation (**Figure 4A**). In contrast, both memory-based conditions produced pronounced changes in cell-wall stiffness and tissue organization, although the resulting patterns differed substantially. In the MT-memory-only condition, the tissue exhibited a heterogeneous distribution of MT orientations. Individual cells developed either longitudinally or transversely aligned MT arrays, leading to corresponding anisotropic patterns of cell-wall stiffening at the cellular level (**Figure 4B**). However, because the preferred orientations varied from cell to cell, no coherent directional organization emerged across the tissue. In the MT-memory-with-external-cue condition, MT organization became coordinated across the entire tissue. Cells aligned their MT arrays along a common orientation determined by the external cue, resulting in a corresponding alignment of cell-wall stiffness patterns throughout the tissue (**Figure 4C).** Consequently, anisotropic reinforcement occurred coherently across neighboring cells rather than independently within individual cells. To quantify the resulting tissue morphology, we measured the aspect ratio (AR) of the tissue over time. Interestingly, the behavior at the tissue scale differed markedly from that observed in single-cell simulations. Both the no-memory and MT-memory-only conditions exhibited largely isotropic tissue growth, with little change in AR over time. In contrast, tissues subjected to MT memory and an external directional cue displayed a progressive increase in anisotropy, reaching approximately 5% higher AR than the no-memory condition (**Figure 4D**). These results suggest that cell-autonomous anisotropic growth is insufficient to generate tissue-scale anisotropy when neighboring cells adopt different orientations. In such cases, local anisotropies are mechanically averaged out through tissue connectivity. However, when MT memory is coordinated by a global external cue, anisotropic growth becomes aligned across cells, allowing individual cellular responses to reinforce one another and collectively generate anisotropic tissue growth.

**Figure 4.**
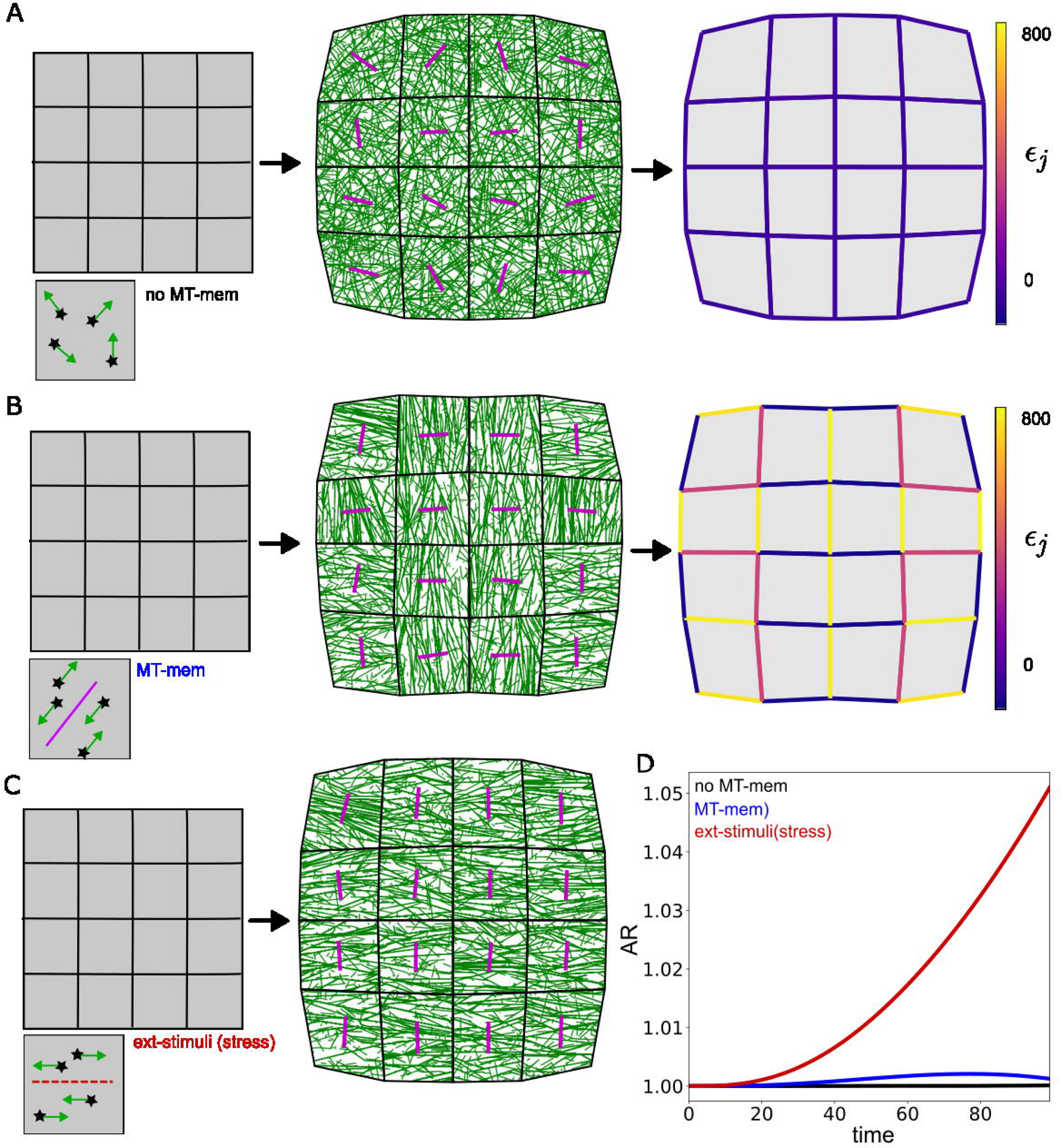
Microtubule memory and external stimulus-based nucleation shape tissue morphogenesis. **(A)** Simulation using isotropic nucleation and default interaction rules: Simulated configuration of MTs on the individual cell surfaces of a tissue initially consisting of rectangular cells, where MTs are simulated using the default interaction rules (as described in Figure 1B). The magenta lines show the orientation of the MT array (left). Heat map, showing wall stiffness at the cell edges (right). **(B)** Simulation using MT memory-based nucleation, where MTs are simulated using the default interaction rules along with the MT memory-based nucleation (as described in Figure 3C). The magenta lines show the orientation of the MT array (top). Heat map, showing wall stiffness at the cell edges (bottom). **(C)** Simulated configuration of MTs on the individual cell surfaces (middle), where MTs are simulated using the default interaction rules along with the external stimulus-based nucleation (as described in Figure 3D). The magenta lines show the orientation of the MT array (top). Heat map, showing wall stiffness at the cell edges (bottom). **(D)** Comparative plot of the aspect ratio of the tissues, when simulated using only default MT-MT interaction rules (black), MT-MT interaction rules along with MT memory-based nucleation (blue), and MT-MT interaction rules along with external-stimuli-based nucleation (green). For vertex model, time unit is nondimensional with step, dt = 0.00001. For details on parameters please refer to **Table 1**.

### MT self-organization and tissue geometry coordinate tissue growth and deformation

Finally, we simulated tissue consisted of a concentric arrangement of cells, with a central octagonal domain representing the meristematic region, surrounded by successive tissue layers corresponding to the cortex, endodermis, epidermis, and other root tissues (**Figure 5A**). This geometry captures key structural features of the Arabidopsis root cross-section and provides a framework for investigating how cortical MT organization influences growth coordination across different tissue layers. Here, also we examined three biologically motivated scenarios: (i) cells lacking persistent MT organization, representing weakly ordered cortical arrays (no MT memory); (ii) cells in which MT organization was influenced by previously established cellulose and MT patterns (MT memory derived from pre-existing MT), and (iii) cells in which MT organization was guided by an external directional signal, mimicking regulation by developmental or mechanical cues (MT memory guided by an external directional cue). To characterize the resulting mechanical states, we quantified the tension along cell edges, defined as *T_j_* = -*ε_j_* (*I_j_* - *I*_0_). The no-memory condition (**Figure 5B**) and the externally guided condition (**Figure 5E**) exhibited similar large-scale mechanical patterns, with positive edge tensions concentrated in the inner tissue layers and negative tensions at the tissue periphery. In contrast, tissues exhibiting inherited MT organization displayed a distinct mechanical state, characterized by near-zero tensions in the outer cell layers (**Figure 5D**). These results suggest that the history of MT organization can influence how mechanical stresses are distributed across the tissue, potentially contributing to the emergence of region-specific growth behaviors during root development.

**Figure 5.**
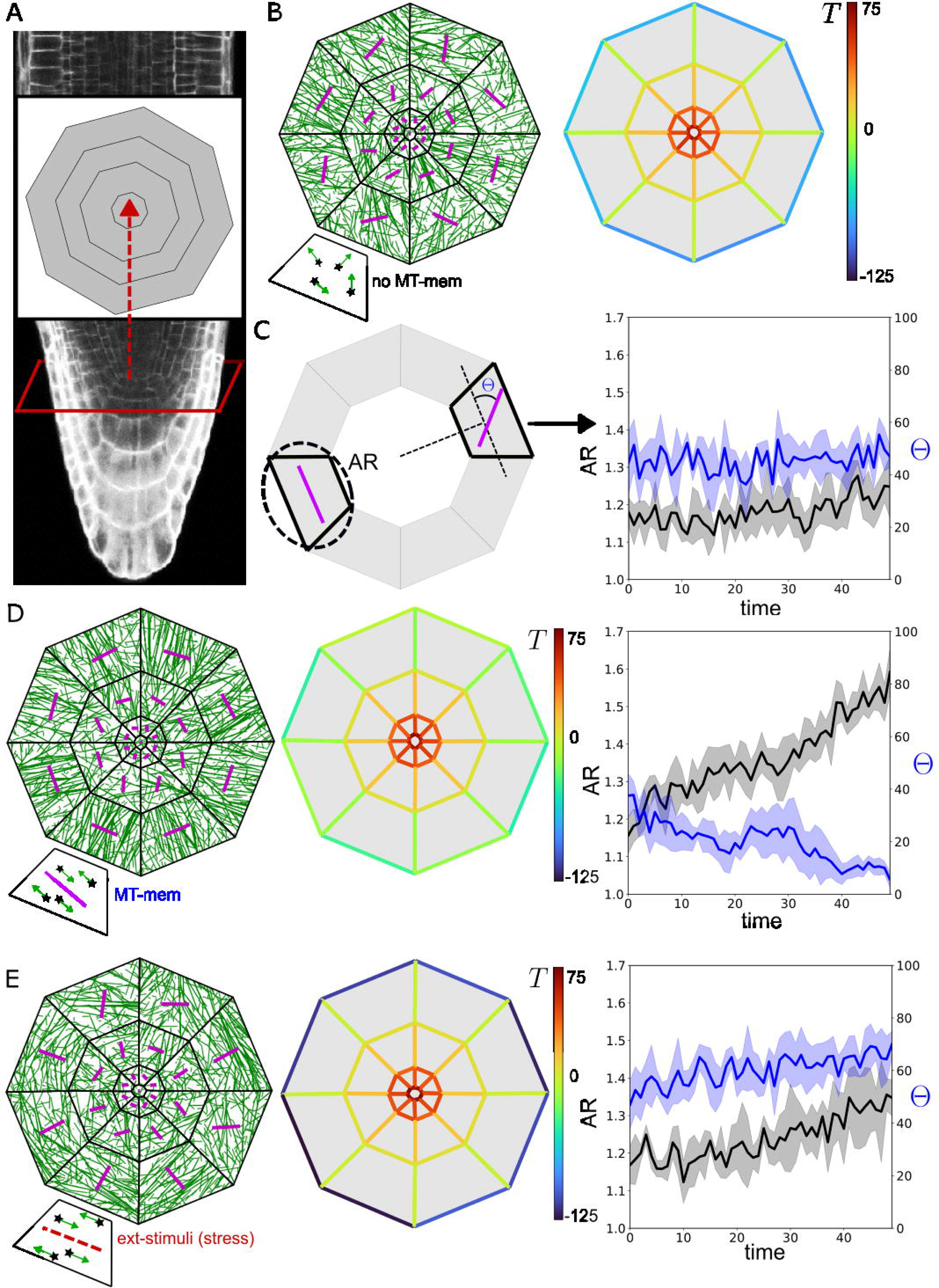
Multiscale simulation of MT dynamics in a biologically realistic root niche tissue model. **(A)** Schematic representation of *Arabidopsis* root tip (bottom, confocal image), where the tissue is consisted of a concentric arrangement of cells, with a central octagonal domain representing the meristematic region, surrounded by successive tissue layers corresponding to the cortex, endodermis, epidermis, and other root tissues. **(B)** Simulation results for the no-MT-memory condition. Left, Tissue geometry with overlaid MT orientation (green lines) and resultant alignment vectors (magenta) for each cell. Right, Spatial distribution of edge tension (T) across the tissue, color-coded according to the scale shown (red: high tension; blue: negative/compressive tension). **(C)** Quantification scheme and baseline (no-memory) dynamics. Left: Schematic illustrating the aspect ratio (AR) measurement (dashed circle) and the MT orientation angle θ, measured relative to the axis perpendicular to the radial (centre-pointing) direction of each cell. Right: Aspect ratio (black, left axis) and θ (blue, right axis) plotted over time for peripheral cells in the no-memory condition; shaded regions indicate variability across the eight peripheral cells. **(D)** Simulated MT configuration through MT-memory-based nucleation condition, shown as in (B), together the heat map showing edge tension at the cell edges (middle) and AR and θ plotted over time (right). MTs at the periphery align longitudinally (parallel to the radial axis), and AR increases progressively over time while corresponding values of θ decreases. **(E)** Simulated MT configuration through MT-memory-with-external-cue condition, shown as in (B) and (D). Heat map, showing edge--tension at the cell edges (middle) and AR and θ plotted over time (right). MTs at the periphery align transversely in response to the applied external stress (dashed red line, inset), with θ remaining close to 90° and comparatively modest change in AR over time. For vertex model, time unit is nondimensional with step, dt = 0.00001. For details on parameters please refer to **Table 1**.

To investigate the relationship between MT organization and cell morphology, we focused on cells in the outermost tissue layer, analogous to epidermal cells that experience strong geometric and mechanical constraints. For each cell, we defined a reference vector tangent to the tissue circumference and measured the angle (Θ) between this direction and the dominant MT-array orientation. Cell shape anisotropy was quantified by fitting an ellipse to each cell and calculating its aspect ratio (AR) (**Figure 5C**). Analysis of Θ and AR revealed a strong coupling between MT organization and cell shape. In tissues where MT organization was inherited from pre-existing MT and cellulose patterns, cortical MT arrays progressively aligned along the circumferential direction of the tissue. This reorientation was accompanied by a decrease in Θ and a corresponding increase in AR over time (**Figure 5D**), indicating preferential cell elongation along the tissue circumference. Such behavior is consistent with experimental observations that cell geometry can influence cortical MT alignment, thereby guiding anisotropic growth. These findings suggest that tissue geometry itself can act as an intrinsic organizational cue that promotes coordinated MT orientation and directional growth. In contrast, when MT organization was influenced by an externally imposed directional cue, MT arrays progressively aligned with the prescribed signal rather than the geometry-derived circumferential orientation. Consequently, Θ increased over time despite continued tissue growth (**Figure 5E**). This result demonstrates that global developmental or mechanical signals can override geometry-driven MT alignment, allowing tissues to adopt growth patterns that are not solely determined by local geometry. Together, these findings suggest that plant tissues may integrate intrinsic geometric information with external regulatory cues to coordinate MT organization and thereby control growth anisotropy during development.

## Discussion

Understanding how subcellular organization gives rise to tissue-scale growth patterns remains a central challenge in plant morphogenesis. This challenge underscores the importance of computational multiscale modeling, particularly because many key mechanical quantities, including tissue stress distributions and force transmission pathways, remain difficult to measure directly in living tissues. In this study, we developed a coupled computational framework that integrates a dynamic vertex model of tissue mechanics with a stochastic model of cortical microtubule (MT) self-organization, thereby linking subcellular cytoskeletal dynamics to emergent tissue-scale growth behavior. The framework incorporates explicit bidirectional feedback in which MT organization determines anisotropic cell-wall stiffness, whereas evolving cell geometry influences subsequent MT alignment. By exploiting the separation of timescales between rapid MT reorganization and slower tissue deformation, the model efficiently captures mechanochemical interactions across subcellular, cellular, and tissue scales. Our simulations demonstrate that differences in MT self-organization alone are sufficient to generate qualitatively distinct growth regimes. Unbiased MT dynamics produce weakly ordered arrays and nearly isotropic tissue expansion, whereas interaction-mediated alignment generates strongly ordered arrays that drive anisotropic growth and tissue elongation. In addition, spatial heterogeneity in MT organization gives rise to mechanically distinct domains exhibiting contrasting growth behaviors. Together, these findings show that stochastic MT self-organization coupled to mechanical feedback constitutes a minimal physical mechanism sufficient to generate tissue-scale growth anisotropy without externally imposed growth orientations or long-range coordinating signals.

A central contribution of this work is the explicit mechanochemical feedback between cytoskeletal organization and tissue mechanics. Unlike continuum models that describe averaged tissue properties, the present framework resolves individual cell geometries together with stochastic MT dynamics, enabling direct investigation of how local cytoskeletal interactions generate coordinated tissue-scale behavior. This feedback provides a plausible mechanism for the amplification and stabilization of growth anisotropy and is consistent with experimental evidence showing that both cell shape and mechanical stress influence MT orientation (Wasteneys & Ambrose, 2009; Zhao et al., 2020). The model further predicts that the transition between isotropic and anisotropic growth is controlled by the strength of MT-guided wall reinforcement, suggesting that modulation of MT alignment, stability, or cellulose deposition may provide an effective mechanism for regulating tissue-scale growth patterns in vivo (Baskin et al., 1999). The emergence of tissues containing both isotropic and anisotropic regions suggests that spatially patterned cytoskeletal organization may be sufficient to establish developmentally distinct growth domains within a single tissue.

Several limitations should be noted. The current model does not include cell division, biochemical signaling pathways such as auxin transport, or fully three-dimensional tissue geometries (Fu & Harberd, 2003). Incorporating these processes, together with quantitative comparisons to experimental measurements of cell shape, MT orientation, and growth rates, will be essential for establishing predictive validity. Future extensions based on experimentally reconstructed cell geometries and more detailed mechanochemical regulation should enable increasingly realistic simulations of developing plant organs. More broadly, this study illustrates how multiscale computational modeling can bridge molecular-scale cytoskeletal dynamics with organ-level morphogenesis. Our results suggest that spatial variation in cytoskeletal organization may represent a fundamental organizing principle underlying developmental pattern formation in plant tissues.

## Acknowledgement

We thank IISER Thiruvananthapuram for providing the research infrastructure and facilities.

## Author Contribution

AKS and AB: Conceptualization, Computational Modelling, Writing - original draft

OP: Writing - original draft.

BC: Conceptualization; Project administration; Writing - review and editing.

## Conflicts of Interest declarations in manuscripts

The authors declare no competing interests.

## Data and Coding Availability Statement

C++ code developed for this project can be made available upon request

## References

1. Alonso Baez, L., Bjørkøy, A., Saffioti, F., Morghen, S., Amanda, D., Tichá, M., Besten, M., Ivanova, A., Sprakel, J., Stokke, B. T., & Hamann, T. (2026). The mechanical properties of Arabidopsis thaliana roots adapt dynamically during development and to stress. Science Advances, 12(8), eaeb0032. 10.1126/sciadv.aeb0032

2. Alt, S., Ganguly, P., & Salbreux, G. (2017). Vertex models: From cell mechanics to tissue morphogenesis. Philosophical Transactions of the Royal Society B: Biological Sciences, 372(1720), 20150520. 10.1098/rstb.2015.0520

3. Ambrose, C., Allard, J. F., Cytrynbaum, E. N., & Wasteneys, G. O. (2011). A CLASP-modulated cell edge barrier mechanism drives cell-wide cortical microtubule organization in Arabidopsis. Nature Communications, 2(1), 430. 10.1038/ncomms1444

4. Baskin, T. I. (2005). ANISOTROPIC EXPANSION OF THE PLANT CELL WALL. Annual Review of Cell and Developmental Biology, 21(Volume 21, 2005), 203–222. 10.1146/annurev.cellbio.20.082503.103053

5. Baskin, T. I., Meekes, H. T. H. M., Liang, B. M., & Sharp, R. E. (1999). Regulation of Growth Anisotropy in Well-Watered and Water-Stressed Maize Roots. II. Role of Cortical Microtubules and Cellulose Microfibrils. Plant Physiology, 119(2), 681–692. 10.1104/pp.119.2.681

6. Beemster, G. T. S., & Baskin, T. I. (1998). Analysis of Cell Division and Elongation Underlying the Developmental Acceleration of Root Growth in Arabidopsis thaliana. Plant Physiology, 116(4), 1515–1526. 10.1104/pp.116.4.1515

7. Bichet, A., Desnos, T., Turner, S., Grandjean, O., & Höfte, H. (2001). BOTERO1 is required for normal orientation of cortical microtubules and anisotropic cell expansion in Arabidopsis. The Plant Journal, 25(2), 137–148. 10.1111/j.1365-313X.2001.00946.x

8. Boudaoud, A. (2010). An introduction to the mechanics of morphogenesis for plant biologists. Trends in Plant Science, 15(6), 353–360. 10.1016/j.tplants.2010.04.002

9. Bozorg, B., Krupinski, P., & Jönsson, H. (2014). Stress and Strain Provide Positional and Directional Cues in Development. PLoS Computational Biology, 10(1), e1003410. 10.1371/journal.pcbi.1003410

10. Chakrabortty, B., Blilou, I., Scheres, B., & Mulder, B. M. (2018). A computational framework for cortical microtubule dynamics in realistically shaped plant cells. PLOS Computational Biology, 14(2), e1005959. 10.1371/journal.pcbi.1005959

11. Dixit, R., & Cyr, R. (2004). Encounters between Dynamic Cortical Microtubules Promote Ordering of the Cortical Array through Angle-Dependent Modifications of Microtubule Behavior. The Plant Cell, 16(12), 3274–3284. 10.1105/tpc.104.026930

12. Dolan, L., Janmaat, K., Willemsen, V., Linstead, P., Poethig, S., Roberts, K., & Scheres, B. (1993). Cellular organisation of the Arabidopsis thaliana root. Development, 119(1), 71–84. 10.1242/dev.119.1.71

13. Eren, E. C., Dixit, R., & Gautam, N. (2015). Stochastic models for plant microtubule self-organization and structure. Journal of Mathematical Biology, 71(6), 1353–1385. 10.1007/s00285-015-0860-9

14. Fu, X., & Harberd, N. P. (2003). Auxin promotes Arabidopsis root growth by modulating gibberellin response. Nature, 421(6924), 740–743. 10.1038/nature01387

15. Green, P. B. (1962). Mechanism for Plant Cellular Morphogenesis. Science, 138(3548), 1404–1405. 10.1126/science.138.3548.1404

16. Hamant, O., & Haswell, E. S. (2017). Life behind the wall: Sensing mechanical cues in plants. BMC Biology, 15(1), 59. 10.1186/s12915-017-0403-5

17. Hamant, O., Heisler, M. G., Jönsson, H., Krupinski, P., Uyttewaal, M., Bokov, P., Corson, F., Sahlin, P., Boudaoud, A., Meyerowitz, E. M., Couder, Y., & Traas, J. (2008). Developmental Patterning by Mechanical Signals in Arabidopsis. Science, 322(5908), 1650–1655. 10.1126/science.1165594

18. Hashimoto, T. (2011). Microtubule and Cell Shape Determination. In B. Liu (Ed.), The Plant Cytoskeleton (pp. 245–257). Springer. 10.1007/978-1-4419-0987-9_11

19. Huang, C., Wang, Z., Quinn, D., Suresh, S., & Hsia, K. J. (2018). Differential growth and shape formation in plant organs. Proceedings of the National Academy of Sciences, 115(49), 12359–12364. 10.1073/pnas.1811296115

20. Li, J., Szymanski, D. B., & Kim, T. (2023). Probing stress-regulated ordering of the plant cortical microtubule array via a computational approach. BMC Plant Biology, 23(1), 308. 10.1186/s12870-023-04252-5

21. Lindeboom, J. J., Lioutas, A., Deinum, E. E., Tindemans, S. H., Ehrhardt, D. W., Emons, A. M. C., Vos, J. W., & Mulder, B. M. (2013). Cortical Microtubule Arrays Are Initiated from a Nonrandom Prepattern Driven by Atypical Microtubule Initiation. Plant Physiology, 161(3), 1189–1201. 10.1104/pp.112.204057

22. Lindeboom, J. J., Nakamura, M., Hibbel, A., Shundyak, K., Gutierrez, R., Ketelaar, T., Emons, A. M. C., Mulder, B. M., Kirik, V., & Ehrhardt, D. W. (2013). A Mechanism for Reorientation of Cortical Microtubule Arrays Driven by Microtubule Severing. Science, 342(6163), 1245533. 10.1126/science.1245533

23. Liu, J., Wang, B., Zhang, Y., Wang, Y., Kong, J., Zhu, L., Yang, X., & Zha, G. (2014). Microtubule dynamics is required for root elongation growth under osmotic stress in Arabidopsis. Plant Growth Regulation, 74(2), 187–192. 10.1007/s10725-014-9910-3

24. Marconi, M., & Wabnik, K. (2021). Shaping the Organ: A Biologist Guide to Quantitative Models of Plant Morphogenesis. Frontiers in Plant Science, 12. 10.3389/fpls.2021.746183

25. Panteris, E., Adamakis, I.-D. S., Daras, G., Hatzopoulos, P., & Rigas, S. (2013). Differential Responsiveness of Cortical Microtubule Orientation to Suppression of Cell Expansion among the Developmental Zones of Arabidopsis thaliana Root Apex. PLOS ONE, 8(12), e82442. 10.1371/journal.pone.0082442

26. Panteris, E., Adamakis, I.-D. S., Daras, G., & Rigas, S. (2015). Cortical microtubule patterning in roots of Arabidopsis thaliana primary cell wall mutants reveals the bidirectional interplay with cell expansion. Plant Signaling & Behavior, 10(6), e1028701. 10.1080/15592324.2015.1028701

27. Panteris, E., Diannelidis, B.-E., & Adamakis, I.-D. S. (2018). Cortical microtubule orientation in Arabidopsis thaliana root meristematic zone depends on cell division and requires severing by katanin. Journal of Biological Research-Thessaloniki, 25(1), 12. 10.1186/s40709-018-0082-6

28. Paredez, A. R., Somerville, C. R., & Ehrhardt, D. W. (2006). Visualization of Cellulose Synthase Demonstrates Functional Association with Microtubules. Science, 312(5779), 1491–1495. 10.1126/science.1126551

29. Sugimoto, K., Williamson, R. E., & Wasteneys, G. O. (2000). New Techniques Enable Comparative Analysis of Microtubule Orientation, Wall Texture, and Growth Rate in Intact Roots of Arabidopsis. Plant Physiology, 124(4), 1493–1506. 10.1104/pp.124.4.1493

30. Wasteneys, G. O., & Ambrose, J. C. (2009). Spatial organization of plant cortical microtubules: Close encounters of the 2D kind. Trends in Cell Biology, 19(2), 62–71. 10.1016/j.tcb.2008.11.004

31. Yu, J., Zhang, Y., & Cosgrove, D. J. (2024). The nonlinear mechanics of highly extensible plant epidermal cell walls. Proceedings of the National Academy of Sciences, 121(2), e2316396121. 10.1073/pnas.2316396121

32. Zhao, F., Du, F., Oliveri, H., Zhou, L., Ali, O., Chen, W., Feng, S., Wang, Q., Lü, S., Long, M., Schneider, R., Sampathkumar, A., Godin, C., Traas, J., & Jiao, Y. (2020). Microtubule-Mediated Wall Anisotropy Contributes to Leaf Blade Flattening. Current Biology, 30(20), 3972–3985.e6. 10.1016/j.cub.2020.07.076

